# Distinct RPA functions promote eukaryotic DNA replication initiation and elongation

**DOI:** 10.1101/2022.09.30.510360

**Authors:** Alexandra M. Pike, Caitlin M. Friend, Stephen P. Bell

## Abstract

Single-stranded DNA binding proteins (SSBs) are essential for DNA replication across all domains of life, but vary significantly in their structure and subunit composition. The eukaryotic SSB, Replication Protein A (RPA), serves critical functions in DNA replication, the DNA damage response, and DNA repair. We sought to determine the requirements for RPA during eukaryotic DNA replication initiation and elongation. To determine whether the ssDNA-binding activity is sufficient, we tested SSBs from different domains of life in reconstituted *S. cerevisiae* origin unwinding and DNA replication reactions. Interestingly, *E. coli* SSB, but not T4 bacteriophage Gp32, fully substitutes for RPA in promoting origin DNA unwinding. Using RPA mutants, we found that only large, multimeric complexes with multiple DNA-binding domains support origin unwinding. In contrast, our studies demonstrated that eukaryotic replication fork function requires specific RPA domains for normal leading- and lagging-strand DNA synthesis. Together, these results reveal new requirements for ssDNA-binding proteins in eukaryotic replication origin unwinding and uncover RPA domains that are critical for faithful replication fork function.

## Introduction

Eukaryotic DNA replication requires the progressive assembly of protein complexes at origins of replication (Bell and Labib, 2016). Origins are licensed during the G1 phase of the cell cycle, when two hexameric Mcm2-7 helicases are loaded onto each origin as a head-to-head double hexamer encircling double-stranded DNA (dsDNA)(Evrin et al., 2009; Gambus et al., 2011; Li et al., 2015; Remus et al., 2009). Upon S-phase entry, S-phase cyclin-dependent kinase (S-CDK) and Dbf4-dependent kinase (DDK) along with additional replication proteins activate the helicase for DNA unwinding (Parker et al., 2017). Once activated, the helicases act separately to bidirectionally unwind DNA (Li and O’Donnell, 2018). The activated helicases and the single-stranded DNA (ssDNA) they generate recruit the remainder of the DNA synthesis machinery to form a bidirectional pair of replication forks(Yao and O’Donnell, 2021).

Helicase activation is the committed step of replication initiation and involves recruitment of key proteins to the helicase and initiation of DNA unwinding. Nine proteins, referred to as activation factors, are sufficient to activate loaded helicases *in vitro(Yeeles* et al., 2015). Eight of the activation factors coordinate the recruitment of Cdc45 and GINS to Mcm2-7 to form the CMG (Cdc45/Mcm2-7/GINS) complex (Bell and Labib, 2016). Cdc45 and GINS directly activate both the ATPase and helicase activity of Mcm2-7(Ilves et al., 2010), converting the CMG complex the active form of the replicative DNA helicase. The final activation factor, Mcm10, acts after CMG formation to further activate the helicase for DNA unwinding (van Deursen et al., 2012; Douglas et al., 2018; Langston and O’Donnell, 2019; Lõoke et al., 2017). Experiments omitting individual activation factors reveal several steps in origin DNA unwinding (Douglas et al., 2018; Lewis et al., 2022). First, CMG formation results in a small amount of DNA melting (~6-7bp per helicase). Addition of Mcm10 stimulates CMG to unwind a limited amount of additional DNA (~10bp). Finally, the eukaryotic single-stranded DNA binding protein (SSB) Replication Protein A (RPA) is required for extensive DNA unwinding (Douglas et al., 2018).

RPA is an essential SSB in eukaryotes that binds ssDNA in a sequence-nonspecific manner (Brill and Stillman, 1991; Wold, 1997; Wold and Kelly, 1988). In addition to its requirement for origin unwinding (Douglas et al., 2018) and stimulation of CMG DNA unwinding (Kose et al., 2020), RPA also protects ssDNA and coordinates DNA damage signaling and DNA repair (Caldwell and Spies, 2020). RPA is heterotrimeric protein that has a conserved domain structure across eukaryotes (Figure 1A)(Wold, 1997). The largest subunit, called Rfa1 in *Saccharomyces cerevisiae,* consists of four oligonucleotide/oligosaccharide-binding (OB)-fold domains, three of which bind DNA (OB-A, OB-B, and OB-C) and another that functions as a protein-interaction domain (OB-F, (Prakash and Borgstahl, 2012). The middle subunit, Rfa2, contributes a fourth DNA-binding OB-fold domain (OB-D), and a C-terminal winged-helix (WH) domain. Rfa3 is the smallest subunit, consisting of a single OB-fold (OB-E) that is required for trimerization of the complex but does not bind DNA. The three proteins form a tight structurally conserved trimerization core between OB-folds C, D, and E (Figure 1A)(Bochkareva et al., 2002).

**Figure 1.**
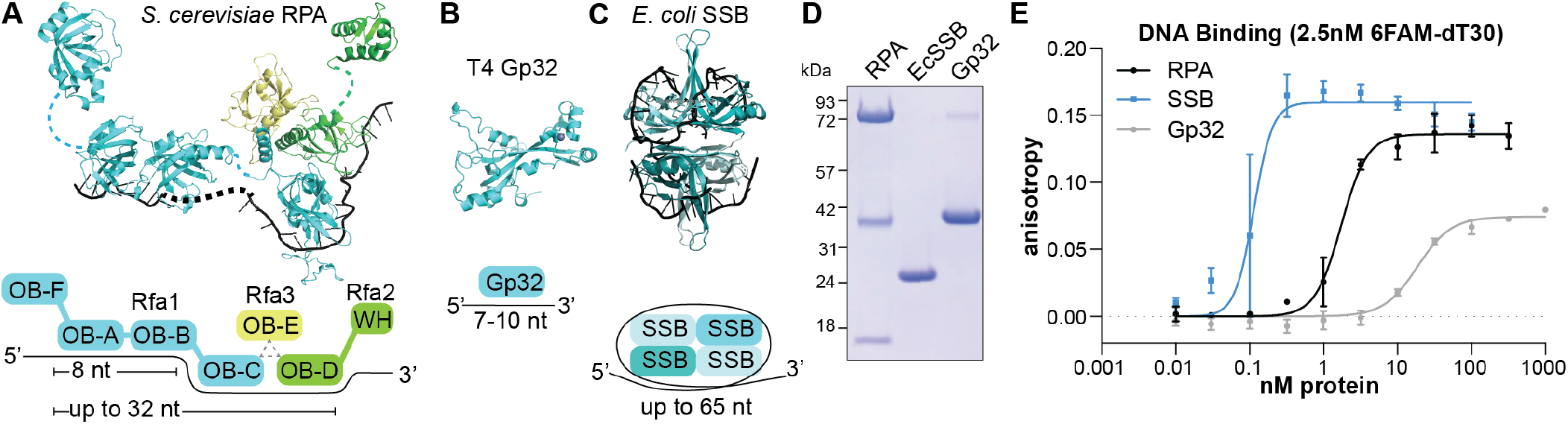
Replicative Single-Stranded Binding proteins vary in their structure and affinity for ssDNA. **A-C.** Structure and cartoon diagram of the subunits and DNA-binding modes of **A.** the RPA heterotrimer, constructed with structures of OB-F (5OMB), OB-A and OB-B bound to ssDNA (1JMC), the OB-C/D/E bound to ssDNA (6I52), and Rfa2-WH (1Z1D), connected by dashed lines representing unresolved protein and DNA sequence (Rfa1, cyan; Rfa2, green; Rfa3, yellow; ssDNA, black). **B**. T4 Gp32 monomer (1GPC), and **C**. E. coli SSB (EcSSB) homotetramer bound to ssDNA (1EYG). OB = oligonucleotide/oligosaccharide-binding fold; WH = winged-helix domain. **D.** Coomassie stained SDS-PAGE gel of purified RPA, EcSSB, and Gp32. **E.** Fluorescence anisotropy measured with 2.5 nM 6-FAM-labeled oligo-dT30 and the indicated concentrations of RPA, EcSSB, or Gp32. Anisotropy values were plotted against the protein concentrations and fit to the Hill Equation.

RPA associates with ssDNA in a highly dynamic manner. These dynamics have been studied both with isolated domains and in the context of the full-length protein(Pokhrel et al., 2019). RPA can adopt multiple ssDNA-binding conformations with varying affinities using its four DNA-binding domains (OB-A through -D; Figure 1A). Depending on the domains involved, RPA can bind as few as 8 nucleotides and as many as 32 nucleotides of ssDNA (Ahmad et al., 2021; Fanning et al., 2006). Binding affinities of RPA and its sub-domains have been reported in the nanomolar to sub-nanomolar range (Caldwell and Spies, 2020). Although cooperative RPA-ssDNA binding was initially observed (Alani et al., 1992), more recent evidence suggests RPA cooperativity only occurs when RPA is phosphorylated during the DNA-damage response(Yates et al., 2018). Structural studies show that the different RPA DNA-binding domains bind ssDNA in different conformations. OB-A and OB-B bind linear stretches of DNA (Bochkarev et al., 1997), whereas, the structure of the trimerization core shows a dramatic bend in the ssDNA as it wraps around the OB-C and OB-D domains (Fan and Pavletich, 2012; Yates et al., 2018). No structure of the full RPA or RPA-DNA complex has been determined, most likely due to the dynamic nature of RPA and its DNA interactions.

SSBs are present in all domains of life as well as a subset of viruses, but these proteins display a wide variety of structures and ssDNA-binding affinities (reviewed in (Marceau, 2012)). The first SSB characterized was Gp32 (Gene 32 protein), which is a monomeric 34 kDa protein that acts during T4 bacteriophage replication (Figure 1B). Gp32 consists of a single OB-fold that can bind up to 10 nucleotides of ssDNA with a Kd of around 10^-8^ M (Kowalczykowski et al., 1981; Rouzina et al., 2005). The canonical bacterial SSB was initially identified in *E. coli* (referred to here as EcSSB), and binds DNA as a homotetramer (Figure 1C). A single EcSSB tetramer has four OB-fold DNA-binding domains that can adopt multiple DNA-binding conformations, binding as many as 65 nucleotides, and has a stronger DNA-binding affinity, with reported dissociation constants in the subnanomolar range (Lohman, 1994; Naufer et al., 2021). Unlike RPA, both Gp32 and EcSSB exhibit strong cooperative DNA binding (Marceau, 2012). Despite their different ssDNA-binding properties, RPA, Gp32, and EcSSB play an essential role during DNA replication in their cognate organism (Marceau, 2012).

Much of our understanding of RPA function during eukaryotic DNA replication comes from studies of simian virus 40 (SV40) DNA replication in human cells. This viral replication fork uses the SV40 large T-antigen (LTag) as the replicative helicase and a subset of the human DNA synthesis machinery to replicate the SV40 genome. Studies of SV40 DNA replication *in vitro* have shown that neither DNA unwinding nor DNA synthesis occur in the absence of RPA (Ishimi et al., 1994; Tsurimoto and Stillman, 1991; Wobbe et al., 1987). Additionally, human RPA has a unique function during SV40 DNA replication, but not DNA unwinding, that cannot be performed by other SSBs, including yeast RPA(Brill and Stillman, 1989; Kenny et al., 1989). Specific interactions between human RPA and the SV40 replication machinery are presumed to mediate the specificity of this function. For example, human but not yeast RPA can bind LTag to promote primosome assembly (Melendy and Stillman, 1993). Further, RPA can stimulate pol-α/primase activity (Braun et al., 1997; Kenny et al., 1989; Weisshart et al., 1998) and promote the pol-α/primase to pol Δ polymerase switch during lagging-strand synthesis (Dornreiter et al., 1992; Waga and Stillman, 1998; Yuzhakov et al., 1999). However, replication-protein interactions with RPA have not been addressed at a fully eukaryotic DNA replication fork.

Here, we investigate the roles of RPA during eukaryotic DNA replication initiation and elongation. Because RPA has not been observed to interact with CMG helicase, we asked if DNA unwinding requires RPA specifically or only the ssDNA-binding activity. To this end, we substituted either Gp32 or EcSSB for RPA in reconstituted origin unwinding assays. Although Gp32 does not support DNA unwinding, EcSSB supports robust origin DNA unwinding. Using a series of RPA mutants, we investigated the properties common to both RPA and EcSSB, and found that neither multiple DNA-binding domains nor high ssDNA-binding affinity was sufficient to facilitate origin DNA unwinding. Finally, we used fully reconstituted replication assays to test whether the ssDNA-binding proteins that supported origin unwinding also function in replication elongation. Unlike origin unwinding, we found that the replication fork requires the function of at least two RPA domains for normal leading- and lagging-strand DNA synthesis.

## Results

### EcSSB, but not Gp32, substitutes for RPA in origin DNA unwinding assays

To study the role of SSBs during eukaryotic DNA replication origin unwinding, we compared purified yeast RPA, *E. coli* SSB (EcSSB), and T4 bacteriophage Gp32 (Figure 1D). We first verified that all three purified proteins were competent for ssDNA-binding activity using fluorescence polarization assays with a fixed concentration of a fluorescently-labeled 30-nucleotide dT oligonucleotide (6FAM-dT_30_). We incubated this oligo with multiple concentrations of each protein and measured DNA binding. To ensure equilibrium binding, all reactions were performed in high-salt conditions (Binz et al., 2006) and incubated for 30 minutes, after which we observed no further changes in anisotropy. Consistent with previously reported measurements (Bastin-Shanower and Brill, 2001; Kowalczykowski et al., 1981; Naufer et al., 2019; Rouzina et al., 2005), Gp32 showed the lowest binding affinity with an apparent Kd of 18.7 nM, followed by RPA (Kd ~ 1.73 nM), then EcSSB (Kd ~ 0.11 nM) (Figure 1E and Supplemental Table 1). The differences in binding-site size, cooperativity, and affinity of each protein for ssDNA all likely contribute to the different binding affinities in this assay. Importantly, the relative binding affinities agree with previous data, and our downstream assays are all performed under saturating concentrations of the respective protein.

Next, we tested each SSB in an *in vitro* origin DNA unwinding assay using purified budding yeast proteins (Douglas et al., 2018). We detected DNA unwinding using a topologically-constrained circular *ARS1* -containing DNA template, onto which we loaded and activated Mcm2-7 in the presence of a topoisomerase (Figure 2A). The topoisomerase relieves supercoiling generated by DNA duplex unwinding. Reactions are terminated by rapid denaturation of the proteins (including topoisomerase) which inactivates and removes them from the DNA. This treatment allows unwound DNA to re-anneal in the absence of topoisomerase, generating negative supercoils that migrate faster in an agarose gel compared to the relaxed plasmid (Figure 2A). The topological changes observed are DDK-dependent, consistent with this kinase being required for assembly of the CMG helicase (Supplemental Figure 1 and Figure 2B-C, lane 5). As previously observed, extensive DNA unwinding required RPA (Figure 2B-C, lanes 3 vs 4).

**Figure 2.**
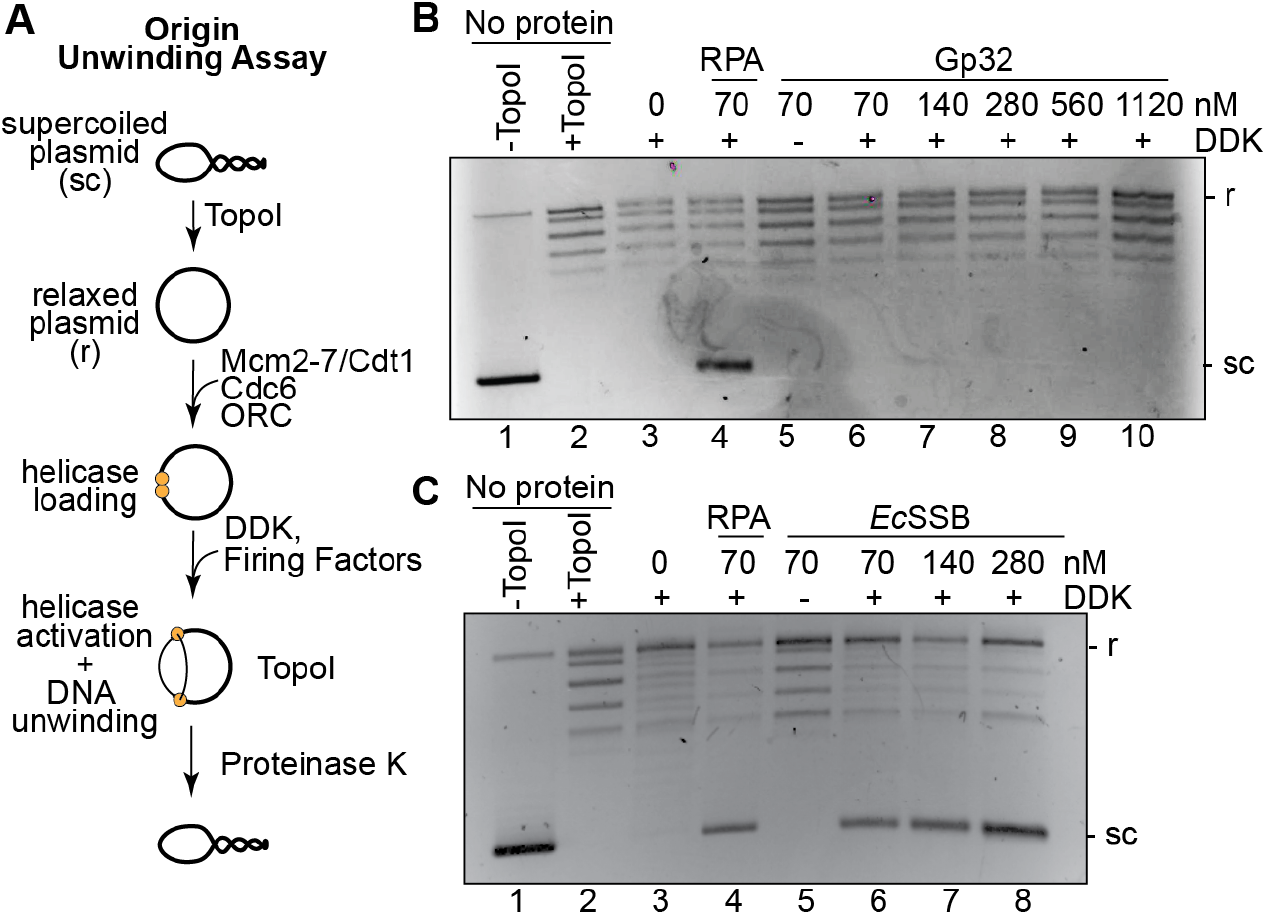
EcSSB, but not Gp32, can support Mcm2-7 unwinding of origin DNA. **A.** Schematic of the origin DNA unwinding assay. **B.** Origin DNA unwinding assays with a two-fold titration series of Gp32 in the absence of RPA. **C.** Origin DNA unwinding assay substituting EcSSB for RPA. In both experiments, “No protein” controls (Lanes 1-2) contain only plasmid +/- Topoisomerase I (Topo I) to show migration of supercoiled (sc) and relaxed (r) plasmids.

If origin unwinding only requires the ssDNA-binding activity of RPA, then other SSBs should support eukaryotic DNA unwinding. However, if additional RPA interactions are required, then other SSBs will fail to substitute for RPA. When we tested Gp32 in the DNA-unwinding assay, we did not observe levels of DNA unwinding above that detected in the absence of any SSB (Figure 2B, lanes 6-10). Increasing the concentration of Gp32 into the micromolar range did not restore DNA unwinding activity, eliminating the possibility that Gp32’s lower ssDNA-binding affinity explains the lack of DNA unwinding (Figure 2B). The inability of Gp32 to support origin unwinding suggests this process requires more than the ssDNA-binding activity of RPA.

Interestingly, when we titrated EcSSB in the unwinding assay, EcSSB supported robust DNA unwinding (Figure 2C, lanes 6-8). We observed similar levels of unwinding between RPA and EcSSB when provided in reactions at the same concentration (Figure 2C, compare lanes 4 and 6). Thus, EcSSB shares the properties of RPA that are required for eukaryotic replication origin unwinding.

### Multiple RPA ssDNA binding domains are required to promote origin unwinding

Gp32 cannot perform at least one RPA function that is essential for eukaryotic DNA unwinding, indicating that more than ssDNA-binding activity is required for this process. Interestingly, EcSSB shares this unknown additional function, implying that it is not a specific interaction with the helicase or an activation factor. One or more of the following shared features of RPA and EcSSB must be required for eukaryotic origin unwinding: multiple ssDNA-binding domains, larger ssDNA-binding footprint, or the ability to dynamically bend or wrap DNA around a multimeric protein complex. Since Gp32 contains only a single DNA-binding domain, we hypothesized that the number of DNA-binding domains could explain the different ability of Gp32 and RPA/EcSSB to facilitate origin unwinding.

To test whether the number of DNA-binding domains and their mode of binding is sufficient to support DNA unwinding, we first generated RPA mutants containing two DNA-binding domains. Structural studies show that RPA and EcSSB both form oligomeric structures with four DNA-a binding domains that can wrap or bend ssDNA (Figure 1A-B), so we tested two different pairs of RPA ssDNA-binding domains that bind DNA in different conformations. RPA^AB^ is composed of OB-A and OB-Bdomains, which are adjacent to one another in Rfa1 and together bind 8 nts of ssDNA in a relatively linear conformation (Bochkarev et al., 1997). In contrast, the RPA trimerization core (RPA^Tri-C^) is a trimer made up OB-domains from three separate subunits (OB-C/D/E; only OB-C and OB-D bind ssDNA) that exerts a local bend in the 19 nt of bound ssDNA (Figure 1A, 3A, (Yates et al., 2018)).

**Figure 3.**
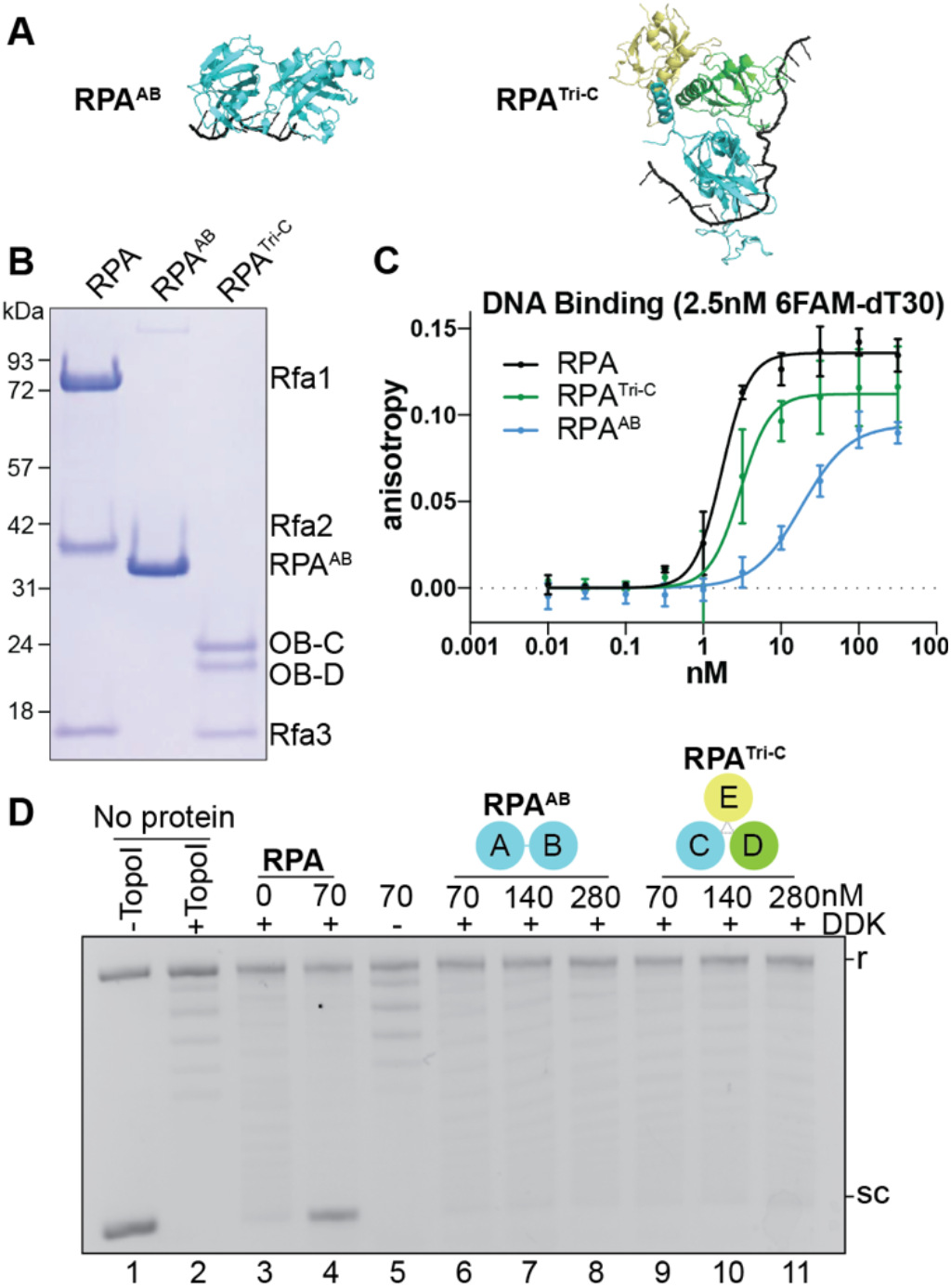
RPA mutants containing two DNA-binding domains do not support unwinding. **A.** Structures of the RPA mutants (1JMC, 6I52). **B.** Coomassie-stained SDS-PAGE gel of RPA, RPA^AB^, and RPATri-c **C.** Fluorescence polarization results of 2.5nM 6-FAM oligo-dT30 incubated with indicated concentrations of RPA^AB^(cyan) or RPA-^Tri-C^ (green). The wild-type RPA results from Figure 1 are included for comparison. **D.** Origin DNA unwinding assays with two-fold titration of RPA^AB^ or RPA^TriC^ (see also Figure 2 legend).

When we tested these RPA mutants in fluorescence polarization assays, both RPA^AB^ and RPA^Tri-C^ displayed lower ssDNA-binding affinity compared to RPA. RPA^AB^ showed ssDNA binding comparable to that of Gp32 (Kd ~ 18.35 nM; Figure 3C), whereas RPA^Tri-C^ DNA-binding affinity was reduced to a lesser extent (Kd ~ 2.97 nM; Figure 3C). When we tested each two-ssDNA-binding-domain SSB in the origin unwinding assay, neither RPA^AB^ nor RPA^Tri-C^ stimulated DNA unwinding. As with Gp32, increasing the concentration of RPA^AB^ or RPA^Tri-C^ to compensate for their lower ssDNA affinity did not restore origin DNA unwinding (Figure 3D and Supplemental Figure 2). We conclude that a ssDNA-binding protein with two DNA-binding domains is not sufficient to promote origin DNA unwinding.

We reasoned that perhaps the high avidity or larger binding site of four DNA-binding domains is required to promote CMG activity. To test this possibility, we made an artificial RPA mutant containing two copies each of the OB-A and OB-B domains in a single polypeptide sequence (RPA^ABAB^), connected by the native linker that separates OB-B from OB-C (Figure 4A-B). Structural evidence shows the OB-A and OB-B domains bind 8nt of ssDNA (Bochkarev et al., 1997), suggesting RPA^ABAB^ can bind at least 16 nt and perhaps more due to the flexible linker between the two AB domains. This RPA mutant showed robust DNA binding, comparable to wild-type RPA (Kd ~ 1.45 nM; Figure 4C). When tested in DNA unwinding assays, however, RPA^ABAB^ failed to promote origin DNA unwinding compared to RPA (Figure 4D, lanes 6-8). These data support the conclusion that neither ssDNA-binding affinity nor number of DNA-binding domains explain the effects of a ssDNA-binding protein on origin unwinding.

**Figure 4.**
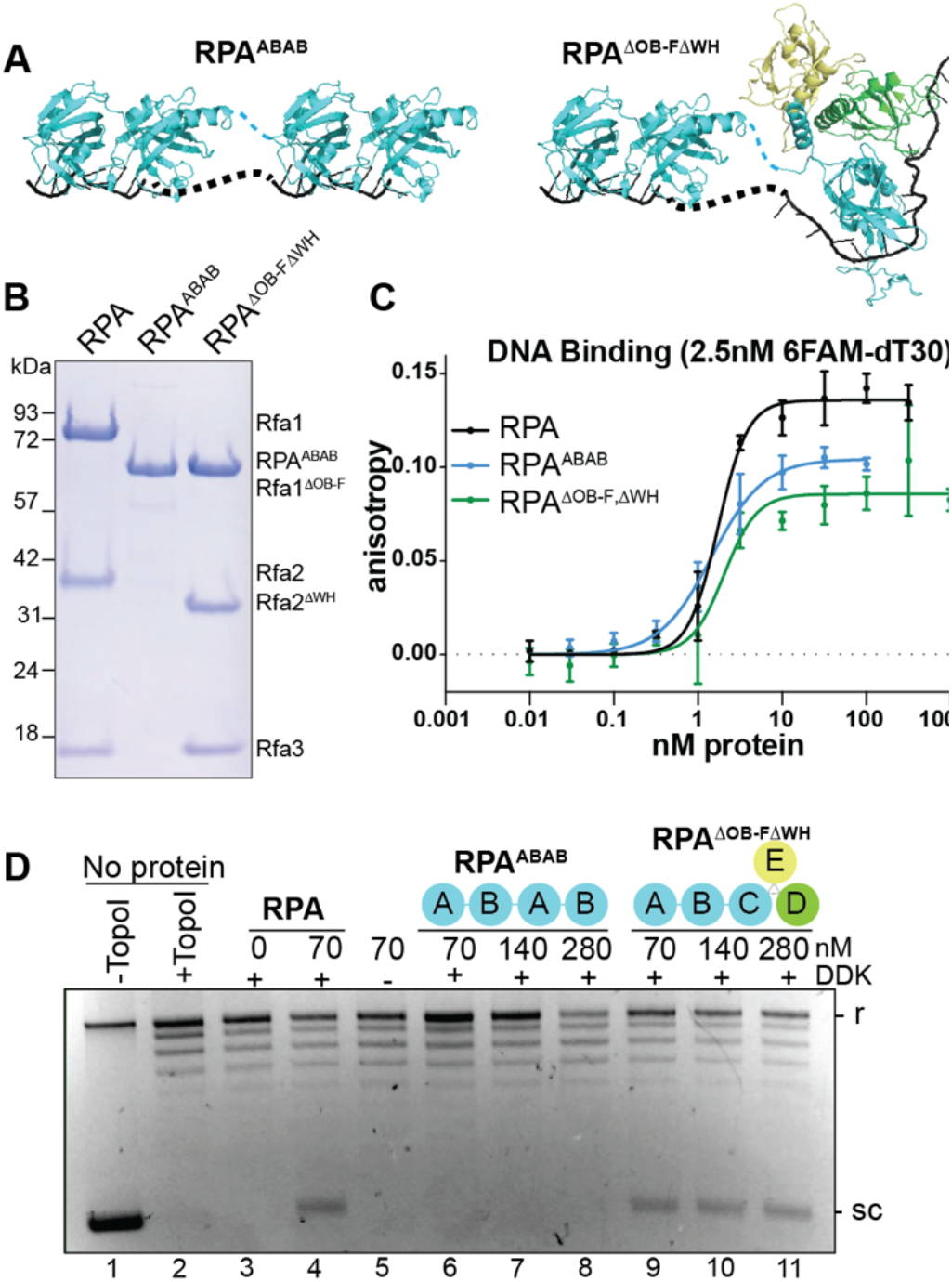
A high-affinity four DNA-binding domain SSB is not sufficient for origin unwinding. **A.** Diagrams of RPA^ABAB^ and RPA^ΔOB-ΔWH^ mutants. **B.** Coomassie-stained SDS-PAGE gel of wildtype and mutant RPA. **C.** Fluorescence polarization results of 2.5 nM 6FAM-oligo-dT30 incubated with indicated concentrations of RPA^ABAB^ or RPA^ΔOB-FΔWH^. The wild-type RPA results from Figure 1 are included for comparison. **D.** Origin DNA unwinding assay with two-fold titration RPA^ABAB^ or RPA^ΔOB-FΔWH^.

To test whether origin unwinding requires the endogenous, multimeric context of RPA, we generated an RPA mutant, RPA^ΔOB-FΔWH^, that contains all four DNA-binding domains but lacks the protein-interaction domains, Rfa1-OB-F and Rfa2-WH (Figure 4A-B). These domains interact with DNA replication and repair proteins but are not involved in DNA binding (Acharya et al., 2021; Collins and Kelly, 1991; Dornreiter et al., 1992; Han et al., 1999; Mer et al., 2000; Prakash and Borgstahl, 2012; Shen et al., 2022; Wold et al., 1989; Zou and Elledge, 2003). Indeed, we found this mutant bound DNA with near wild-type affinity (Figure 4C). In contrast with the RPA^ABAB^ mutant, however, RPA^ΔOB-FΔWH^ supported robust DNA unwinding at all concentrations tested (Figure 4D, lanes 4 vs 9-11). Together, these RPA mutant results indicate that origin unwinding requires the full, multimeric context of RPA ssDNA binding rather than the binding footprint or number of DNA-binding domains alone.

### Replication elongation requires the RPA OB-F and WH domains

We next asked whether ssDNA-binding proteins that support origin unwinding also support DNA synthesis. To monitor replication initiation and elongation, we utilized an *in vitro* reconstituted DNA replication assay on linearized ARS1-containing plasmid DNA (Figure 5A, (Lõoke et al., 2017; Yeeles et al., 2015)). Nascent DNA strands are monitored by incorporation of radiolabeled dNTPs followed by separation by alkaline gel electrophoresis. Omission of Fen1 and DNA ligase from the reactions allows distinction of long leading-strand synthesis products (3000-6000 nt) from shorter lagging-strand products (100-500 nt) (Figure 5, Yeeles et al., 2015). When we tested each of the SSBs that supported in the origin unwinding in the *in vitro* replication assay, we found that DNA synthesis has distinct SSB requirements compared to DNA unwinding.

**Figure 5.**
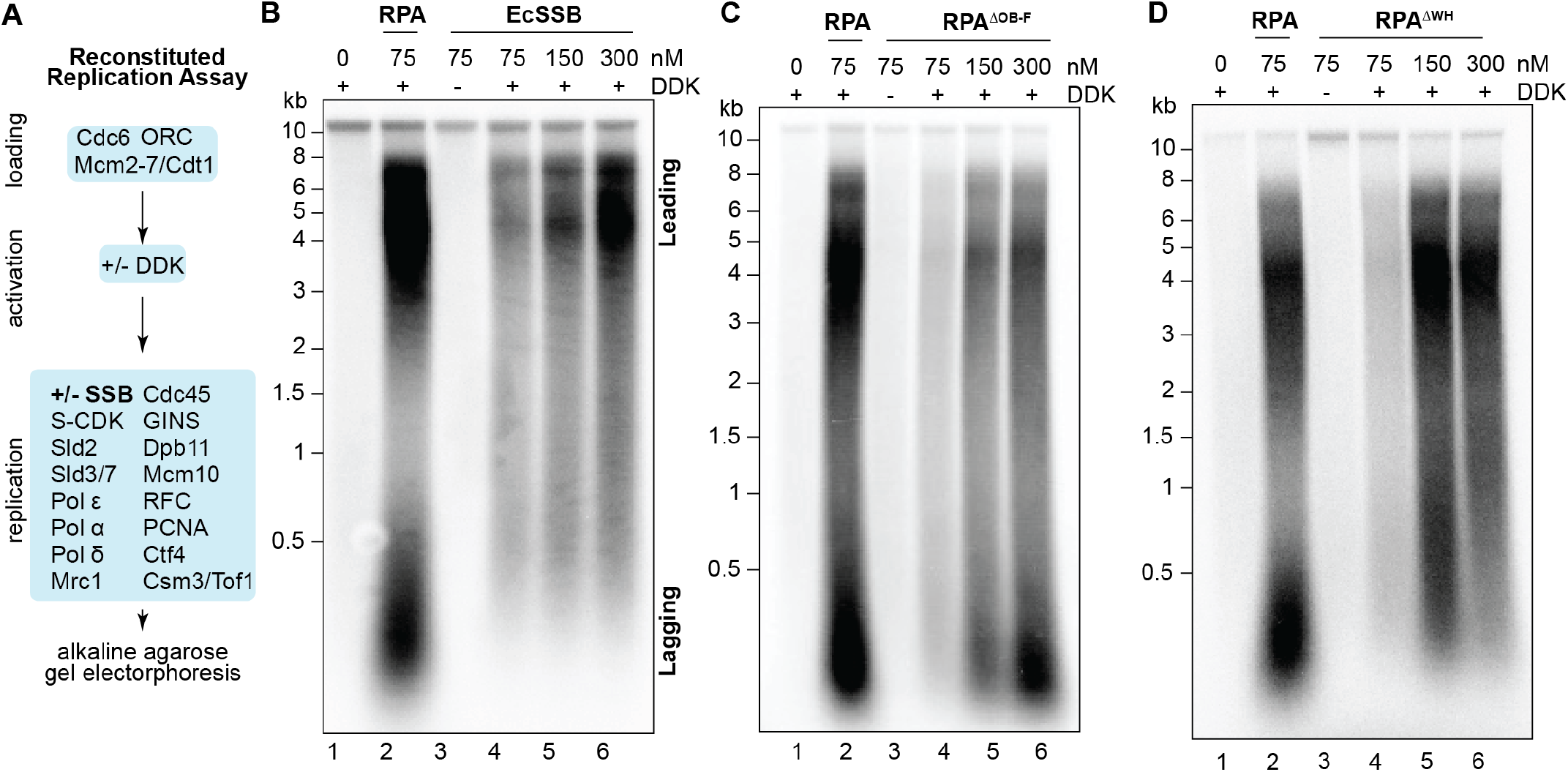
RPA OB-F and WH domains control replication elongation. **A.** Schematic of the reconstituted DNA replication assay. **B-D.** *In vitro* replication with a two-fold titration series of (**B**) EcSSB, (**C**) RPA^ΔOB-F^, or (**D**) RPA^ΔWH^ substituting for RPA. Observed size distribution of normal leading and lagging-strand products is labeled in B. As with the origin DNA unwinding assay, replication is dependent on RPA (compare lanes 1 and 2) and on DDK (compare lanes 2 and 3).

Consistent with the requirement for RPA in unwinding, we saw no DNA replication in absence of any added SSB, but addition of RPA led to robust, DDK-dependent leading- and lagging-strand synthesis (Figure 5, lanes 1-3). Likewise, Gp32 did not support any DNA synthesis (Supplemental Figure 3A). When EcSSB was substituted for RPA in this assay, however, we observed a distinctly different pattern of replication products. When present at the equivalent concentration of RPA (75 nM), EcSSB supported weak DNA-synthesis activity (Figure 5B, lane 4). Thus, while this EcSSB concentration supports robust DNA unwinding (Figure 2), there is a strong defect in initiating and elongating nascent DNA. Increasing EcSSB concentration resulted in a corresponding increase in DNA synthesis, however, the distribution of leading- and lagging-strand products was strikingly different than that observed with RPA. With EcSSB, we consistently observed a much larger fraction of long products (Figure 5B, lanes 5-6), suggesting that lagging-strand DNA synthesis is defective. This could be due to fewer lagging-strand synthesis products or to longer Okazaki fragments that comigrate with the leading strand products. In either scenario, EcSSB fails to promote one or more essential functions at the eukaryotic replication fork.

The failure of EcSSB to support normal leading- and lagging-strand replication suggests that replication requires direct contacts with RPA and the replisome. Interestingly, when we tested the RPA^ΔOB-FΔWH^ mutant that lacks both of RPA’s protein-interaction domains and binds DNA with wild-type affinity (Figure 4), we observed very similar results (Supplemental Figure 3B). This result suggests the OB-F and WH domains control important aspects of DNA replication. To investigate this connection further, we generated deletion mutants lacking either the OB-F or the WH domain. These mutants both support robust origin unwinding at all concentrations tested (Supplemental Figure 3C-D). Strikingly, both RPA^ΔOB-F^ and RPA^ΔWH^ had replication initiation defects when supplied at the same concentration as RPA (Figure 5C-D, lanes 2 vs 4).

Increasing the concentration of either mutant protein increased nucleotide incorporation but resulted in distinctly different patterns of leading- and lagging-strand replication products. Higher concentrations of RPA^ΔOB-F^ led to an accumulation of short products comigrating with lagging-strand products (Figure 5C, lanes 4-6). In contrast, increasing the amount of RPA^ΔWH^ produced replication products that lacked the shorter Okazaki fragments, similar to reactions with EcSSB or with RPA^ΔOB-FΔWH^ (Figure 5D, lanes 4-6). These results show that these two protein-interaction domains controls separate activities during DNA replication elongation.

## Discussion

By substituting other SSBs or mutant versions of RPA, we have gained important insights into the function of RPA during eukaryotic replication initiation and elongation. We found that RPA uses two protein-interaction domains, OB-F and WH, to coordinate DNA synthesis at the eukaryotic replication fork (Figure 5). Interestingly, these domains are dispensable for origin unwinding. Instead, we find that the particular arrangement of ssDNA binding motifs found in RPA and EcSSB but not Gp32 are required for this event.

### Origin DNA remodeling during helicase activation

Our results indicate that preventing strand re-annealing is not the sole purpose of RPA during origin unwinding (Figures 2–4). There are multiple steps of origin DNA unwinding that could require RPA. CMG activation requires remodeling of the helicase and the associated DNA, as the helicase must transition from encircling dsDNA to encircling the leading-strand template ssDNA while excluding the lagging-strand template. CMG formation, which does not require RPA, melts a small amount of the double-stranded DNA within the helicase central pore (Figure 6A)(Lewis et al., 2022), and Mcm10 promotes further duplex melting, observed as a total of two helical turns of DNA per helicase (~20 bp and Figure 6B, top)(Douglas et al., 2018). Because each CMG encircles ~30 bp of DNA, it is unlikely that the non-translocating ssDNA has been extruded entirely from the Mcm2-7 complex in the absence of RPA. This raises the interesting possibility that RPA binding to the melted DNA is required for complete strand extrusion (Figure 6B). Subsequently, the two helicases must pass each other by translocating on opposite ssDNA strands (Figure 6C). Binding of RPA to the non-translocating strand could also stimulate this separation. Finally, RPA binding to the excluded strand stimulates the rate of the CMG helicase unwinding (Kose et al., 2020) and this function could be required to allow enough DNA unwinding to be detected in the topological assay used here.

**Figure 6.**
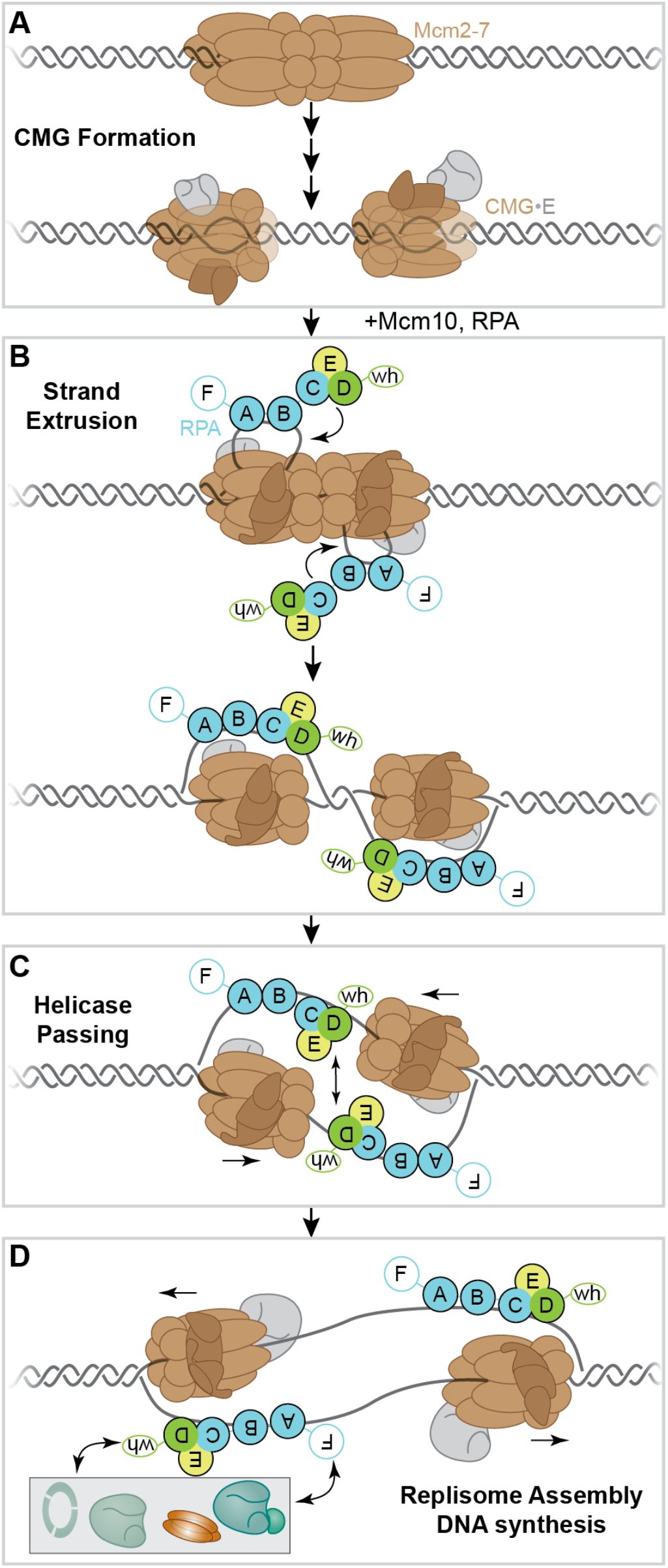
Model for RPA function during replication. **A.** CMG formation begins duplex DNA melting, independent of RPA. **B.** Mcm10 initiates further duplex melting and potentially strand extrusion (illustrated). Transition between different RPA ssDNA-binding modes could promote completion of ssDNA extrusion. **C.** The two activated helicases must pass each other on opposite ssDNA strands at the replication origin. RPA size could facilitate or stabilize strand separation. **D.** The RPA OB-F and WH domains regulate distinct steps during replisome assembly and/or DNA synthesis, most likely by interacting with one or more replicative DNA polymerase or accessory factor.

A role of RPA during strand extrusion could explain the SSB requirements observed in our studies. RPA could promote strand extrusion by engaging with melted DNA in its 8-nt binding mode, then engaging the rest of its DNA-binding domains to convert to the longer ssDNA-binding mode (Figure 6B). This expanding ssDNA binding could pull the remaining ssDNA from the CMG central channel and complete the strand extrusion process. This function could also explain the ability of EcSSB to substitute for RPA, because EcSSB also has both short and long ssDNA-binding modes, depending on how many DNA-binding domains are engaged with ssDNA (Lohman, 1994). RPA effects on ssDNA conformation also could promote strand extrusion. Structural evidence shows that full-length RPA can exert a local bend in the ssDNA (Fan and Pavletich, 2012; Yates et al., 2018). We aimed to test the requirement of this bending using the RPA^Tri-C^ mutant, however, the isolated trimerization core alone may not exert this conformational change. Because the structural data supporting bending by RPA^Tri-C^ was obtained with the full RPA complex, it is possible that the observed bend requires the full complex, even though only the RPA^Tri-C^ domains are resolved. EcSSB also bends ssDNA, wrapping it around the DNA-binding domains (Figure 1, Raghunathan et al., 2000). Like a transition from short to long regions of ssDNA binding, the ability to bend the ssDNA could generate a force that promotes strand extrusion. Importantly, neither multimodal ssDNA binding or DNA bending are known activities of Gp32.

RPA could also promote helicase passing at the replication origin (Figure 6C). Because the Mcm2-7 helicases are loaded in a head-to-head conformation (Evrin et al., 2009; Remus et al., 2009), they must pass each other on opposite strands of DNA before unwinding DNA bidirectionally from the replication origin. RPA could promote this process by eliminating steric dsDNA barriers to this essential step of origin activation. RPA and EcSSB (116 and 75 kDa, respectively) are significantly larger than Gp32 (34 kDa) and contain a clustered OB-fold domain structure that is absent in Gp32 (Figure 1). Although our RPA^ABAB^ mutant approached the size and number of DNA-binding domains of RPA, its three-dimensional structure and DNA binding mode are unlikely to mimic that of RPA or EcSSB. A single RPA DNA-binding OB-fold measures approximately 20-30 Å across, while the Tri-C region of RPA and the diameter of EcSSB each measure approximately 50-70 Å. Such larger structures may serve as a lever to stabilize ssDNA and create the space needed for the large CMG helicases to pass each other on opposite strands of DNA at replication origins.

Finally, RPA could also stimulate CMG processivity on ssDNA. Although RPA modulates the activity of other helicases through direct interactions (Acharya et al., 2021; Shorrocks et al., 2021), the ability of EcSSB and RPA^ΔOB-FΔWH^ to substitute for RPA suggests a direct protein-protein interaction between RPA and the helicase are not required. Instead, we consider that RPA stimulates CMG by interacting with the excluded strand. Structural studies of the CMG complex found that the excluded strand interacts with the helicase central pore (Eickhoff et al., 2019; Georgescu et al., 2017; Goswami et al., 2018). Instead of facilitating DNA unwinding, recent evidence suggests this interaction leads to CMG stalling (Kose et al., 2020). Indeed, RPA binding to the excluded strand stimulates the rate of the CMG helicase (Kose et al., 2020). Therefore, RPA stimulation of the processive CMG helicase activity likely also contributes to the unwinding that we detect in our reconstituted assay. Importantly, these scenarios are not mutually exclusive, thus, it is possible that RPA performs one or more of these functions.

### Replication requires the RPA OB-F and WH domains

Although EcSSB can substitute for RPA in origin unwinding assays, it did not complement RPA function at the replication fork. When provided at the same concentration as RPA, EcSSB and RPA^ΔOB-FΔWH^ displayed decreased signal in reconstituted replication assays, suggesting defects in initiation (Figure 5). At higher concentrations, more DNA synthesis occurred, but with defective leading- and lagging-strand product distribution. Previous work observed that EcSSB could support only rolling-circle replication on a circular DNA template (Devbhandari et al., 2017). Using a defective Pol ε enzyme, the authors showed that rolling-circle replication products were not produced in RPA-containing reactions, suggesting those observed in EcSSB-containing reactions are also the result of Pol ε synthesis. This suggests that perhaps the defects observed in our reactions on a linear template are due to the inability of EcSSB to recruit Pol-α/primase and Pol Δ to replication forks. Our results support the conclusion that direct interactions between RPA and the eukaryotic replisome are required for appropriate replication fork function. SV40 replication experiments also show specific requirements for human RPA at the SV40 replication fork, as neither EcSSB nor yeast RPA can efficiently complement RPA function (Brill and Stillman, 1989; Melendy and Stillman, 1993).

Using individual domain deletions, we found that the Rfa1 OB-F and the Rfa2-WH domains have different functions during replication (Figure 5). The double deletion mutant resembled reactions with EcSSB, with a distribution of larger products but no short Okazaki fragments. Interestingly, the individual deletion of the WH domain recapitulated the results observed with the double deletion. In contrast, deletion of the OB-F domain yielded an opposite effect, resulting in an accumulation of short replication products. These observations suggest that the event stimulated by the WH domain is required before the event that requires the OB-F domain. These domains were chosen for elimination because they are known protein-interaction domains (Caldwell and Spies, 2020), and these results suggest that each domain has a specific interaction that regulates a different step(s) of nascent strand initiation or elongation. Indeed, RPA has been shown to stimulate Pol α/primase activity as well as Pol-α/primase to Pol Δ “polymerase switching” during lagging-strand synthesis (Dornreiter et al., 1992; Waga and Stillman, 1998; Yuzhakov et al., 1999). These activities are candidates for the mechanism through which RPA orchestrates DNA synthesis during replication. Further, since Pol-α/primase initiates both the leading and lagging strands, defective Pol-α/primase regulation would result in the observed reduced replication efficiency observed in these mutants. While Rfa1 and the OB-F domain have been implicated in interactions with Pol-α/primase and RFC, replication fork proteins are not among the proteins known to interact with the Rfa2-WH domain (Caldwell and Spies, 2020). Future studies involving assays for specific steps of replication initiation will be required to determine the precise mechanism by which the OB-F and WH domains regulate replication fork function.

## Acknowledgements

We are grateful to members of the Bell Laboratory for useful discussions. We thank Christian Ramsoomair and Anusha Keerthi for preparation of a subset of the proteins used in this manuscript. We thanks Annie Zhang for comments on the figures and manuscript. S.P.B. is an Investigator of the Howard Hughes Medical Institute. This paper was typeset with the bioRxiv word template by @Chrelli: www.github.com/chrelli/bioRxiv-word-tem-plate

## Author contributions

C.M.F. made the initial observations concerning the function of Gp32 and EcSSB in the origin unwinding and replication assays. A.M.P. generated the RPA mutants and performed all the experiments in the manuscript. A.M.P. and S.P.B wrote the manuscript with input from C.M.F. S.P.B. directed the studies.

## Materials and Methods

### RPA Expression Plasmids and Strains

RPA was expressed and purified from yAE31(Yeeles et al., 2015). RPA protein-interaction domain mutants were made by mutating the respective expression plasmids to generate pRS303-Gal1,10-CBP-Rfa1ΔOB-F/Gal4 and pRS306-Gal1,10-3xFlag-Rfa2ΔWH/Rfa3. The respective plasmids were integrated into yRH101 to generate yAP05 (CBP-Rfa1ΔOB-F, Rfa2, Rfa3); yAP17 (CBP-Rfa1, 3x-Flag-Rfa2ΔWH, Rfa3); yAP18 (CBP-Rfa1ΔOB-F, 3x-Flag-Rfa2ΔWH, Rfa3). Other RPA mutants were expressed in bacteria. To make RPA^AB^, the coding sequence for CBP-OB-A-OB-B was ordered as a gBlock (IDT) and then cloned into the pET3aTr backbone for expression in *E. coli.* RPA^ABAB^ was generated by PCR amplifying OB-A through the N-terminal boundary of OB-C (OB-AB+linker) followed by Gibson assembly into the RPA^AB^ plasmid. To express the trimerization core, the p11d-tscRPA-30MxeHis6 plasmid (Gibb et al., 2014) was modified by truncating the coding sequences for Rfa1 and Rfa2 to contain only OB-C and OB-D, respectively. The resulting plasmid contains an inducible Rfa1-OB-C, Rfa2-OB-D, and Rfa3 coding sequences with an intein, chitin-binding domain, and a 6xHis tag at the Rfa2 C-terminus. All mutations were confirmed by Sanger Sequencing. Additional details on plasmids and yeast strains can be found in Supplementary Tables 2 and 3.

### Protein Expression and Purification

Purified *Escherichia coli* SSB (Sigma-Aldrich S3917) and T4 Gene 32 Protein (Gp32; Roche 10972983001) were obtained from commercial vendors. *Saccharomyces cerevisiae* RPA, RPA^ΔOB-F^, RPA^ΔWH^, and RPA^ΔOB-FΔWH^ were purified from yeast and RPA^AB^, RPA^ABAB^, and RPA^Tri-C^ were purified from *E. coli* as described below.

Wild-type RPA (CBP-Rfa1, Rfa2, and Rfa3) was expressed and purified from yAE31 using calmodulin and HiTrap Heparin columns as described in (Yeeles et al., 2015). RPA^ΔOB-F^ was expressed and purified from yAP05 using the same protocol. Briefly, 8 L of logarithmic culture were alpha-factor arrested and RPA expression was induced with galactose for 3.5 hours. Cells were harvested and ground into powder which was then thawed into Buffer C (25 mM Tris HCl pH 7.2, 10% glycerol, 1mM DTT) with 500 mM NaCl. Lysate was clarified by ultracentrifugation (45 krpm for 90 minutes), then supplemented with 2 mM CaCl2 and bound to a 1 ml calmodulin-affinity column. RPA was eluted with Buffer C supplemented with 200 mM NaCl, 2 mM EDTA, and 2 mM EGTA. RPA-containing fractions were pooled, dialyzed against Buffer C with 50 mM NaCl and 1 mM EDTA, and applied to a 1 ml HiTrap Heparin column equilibrated in the same buffer. RPA was eluted with a 30 column volume (CV) gradient from 50 mm to 1M NaCl in Buffer C + 1mM EDTA. RPA-containing fractions were collected, snap-frozen on liquid nitrogen, and stored at −80°C until use.

For purification of RPA^ΔWH^, lysate from 8L of yAP17 (CBP-Rfa1, 3xFlag-Rfa2-ΔWH, and Rfa3) was prepared as described for RPA and then bound to 1ml Flag resin (Sigma) equilibrated in Buffer C with 100 mM NaCl, washed, and eluted with Buffer C supplemented with 100 mM NaCl and 0.3 mg/ml 3xFlag peptide. RPA-containing fractions were pooled, concentrated, and applied to S200 Increase 10/300 equilibrated in Buffer C with 200 mM NaCl and 1 mM EDTA. Stoichiometric RPA complexes were snap frozen and stored at −80. RPA^ΔOB-FΔWH^ was expressed and purified from 16 L of yAP18 using the approach for WT RPA, except a flag affinity step (as described for RPA^ΔWH^) was added between the calmodulin and heparin columns to enrich for complexes that contained both mutant subunits.

For RPA^AB^ and RPA^ABAB^, the appropriate expression plasmid was transformed into Rosetta(DE3)pLysS cells, expanded to 2 l and induced with 1 mM IPTG during mid-log phase and incubated at 16°C overnight. Cells were pelleted and lysed by sonication in C/500 (25 mM Tris HCl pH 7.2, 10% glycerol, 1mM DTT, 500mM NaCl, and 1mM EDTA). Lysates were then purified by calmodulin and heparin columns as described for WT RPA.

For RPA^Tri-C^, p11d-scTriC was transformed into Rosetta(DE3)pLysS and expressed and purified as in (Lõoke et al., 2017), except after the Ni-NTA and chitin columns, the eluate, which had a significant excess of Rfa2, was pooled and passed over a heparin column. The monomeric Rfa2 saturated the heparin column and Tri-C was concentrated in the flow-through. The flow-through was collected and Tri-C was further purified by binding to a 1 ml MonoQ column equilibrated in Buffer C with 50mM NaCl and 1mM EDTA, then eluted with a 30 CV gradient from 50 mM to 1 M NaCl. Fractions containing trimeric RPA were further purified further by size exclusion on S75 increase in Buffer C with 200 mM NaCl and 1 mM EDTA, snap-frozen, and stored at −80°C.

For reconstituted plasmid unwinding and replication assays, Mcm2-7/Cdt1 and ORC complexes were purified as described previously (Kang et al., 2014). Cdc6 was purified as described in (Frigola et al., 2013). DDK, S-CDK, Sld3/7, Sld2, Dpb11, GINS, Mcm10, and Pol ε were purified as described in (Lõoke et al., 2017) and Cdc45 was purified according to (Posse et al., 2021).

### Fluorescence Polarization Assays

Serial 2-log dilutions of the respective SSB were prepared in 30 mM HEPES pH 7.5, 30 mM NaCl, 0.25 mM EDTA, 10% glycerol, 0.01% NP-40, 1 mM DTT. Proteins were then mixed with 6-FAM-labeled oligo-dT30 (IDT) to a final concentration of 2.5 nM supplemented with 0.5 M NaCl (Binz et al., 2006). The protein-DNA mix was incubated for at least 30 minutes at room temperature to reach equilibrium (Jarmoskaite et al., 2020). Three technical replicates were plated in a black 384-well nonbinding microplate (Greiner Bio-One) and read in a SpectraMax ID5 plate reader. Anisotropy values from the technical replicates were averaged and corrected by subtracting the value from a buffer control that contained no protein. Values from three independent experiments were plotted in GraphPad PRISM and fit to the Hill equation.

### Unwinding Assay

Unwinding assays were performed on 3.8 kb pUC19-ARS1 plasmids as described (Douglas et al., 2018) with modifications. Briefly, 25 fmol of plasmid DNA was relaxed with 0.4 pmol Topo I for 30 min at 30 °C. Then Mcm2-7 loading was performed at 25°C by adding 45 nM ORC, 45 nM Cdc6, and 100 nM Mcm2-7/Cdt1 in 25 mM HEPES-KOH (pH 7.6), 10 mM magnesium acetate, 225 mM potassium glutamate, 2 mM DTT, 0.02% NP-40, 5% glycerol, 5 mM ATP, 20 mM phosphocreatine, and 0.2 μg of creatine kinase for a total volume of 10 μl. After a 25 minute incubation, 1.3 pmol of DDK was added and incubation was continued for a further 30 min. DNA unwinding was then initiated by adding 20 μl of firing-factor mix (0.6 pmol CDK, 1 pmol Sld3/7, 1 pmol Cdc45, 1.24 pmol Sld2, 0.8 pmol Dpb11, 5 pmol GINS, 0.06 pmol Mcm10, 0.6 pmol Pol ε, 0.4 pmol of Topo I, and the indicated SSB amount) in 25 mM HEPES-KOH (pH 7.6), 10 mM magnesium acetate, 250 mM potassium glutamate, 1 mM DTT, 0.02% NP-40, 8% glycerol, 5 mM ATP, and 0.4 mg/ml BSA. After a 40 minute incubation at 25°C, the reaction was quenched with EDTA, Proteinase K, and SDS as described (Douglas et al., 2018). Samples were purified by phenol:chloroform extraction, ethanol precipitation, and run on 1.5% agarose TAE gels at 1.5 V/cm for 16-20 hours before staining with ethidium bromide and imaging.

### Replication Assay

Replication assays were performed on a soluble 11.9kb *ARS1*-containing DNA template that was linearized with NotI and purified by phenol:chloroform extraction and ethanol precipitation. Helicase loading was performed by incubating 10 μl reactions with 0.125 pmol DNA, 0.5 pmol ORC, 0.5 pmol Cdc6, and 1.25 pmol Mcm2-7/Cdt1 in a buffer containing 25 mM HEPES (pH 7.6), 10 mM MgOAc, 2 mM DTT, 100 mM KGlut, 20 mM phosphocreatine (PC), 5 mM ATP, 0.01% NP-40, 5% glycerol, and 0.2 μg of creatine kinase (CK) for 25 minutes at 25°C with shaking on an Eppendorf Thermomixer at 1250 rpm. Then, 1.3 pmol DDK was added and the reaction was incubated for another 25 minutes at 25°C at 1250rpm. After DDK phosphorylation, replication was initiated by adding 20 μl of a replication mix containing 1 pmol CDK, 1 pmol Sld3/7, 2.6 pmol Cdc45, 1.24 pmol Sld2, 0.8 pmol Dpb11, 5 pmol GINS, 0.02 pmol Mcm10, 0.6 pmol Pol ε, 2 pmol pol-α/primase, 0.6 pmol Ctf4, 0.5 pmol RFC, 0.4 pmol PCNA, 0.5 pmol Mrc1, 0.6 pmol Csm3/Tof1, 0.2 pmol Pol Δ, and the indicated SSB concentration in replication buffer (12.5 mM HEPES-KOH (pH 7.6), 5 mM magnesium acetate, 125 mM potassium glutamate, 1 mM DTT, 0.01% NP-40, 4% glycerol, 1.5 mM ATP, 10 mM phosphocreatine, 3 μg of creatine kinase, 0.2 mg/ml BSA, 100 μM rNTP, 10 μM dNTP, and 10 μCi [α-P^32^]-dCTP). After a 60-minute incubation at 25°C while shaking at 1250 rpm, reactions were quenched with EDTA and unincorporated nucleotides were removed with Illustra MicroSpin G-50 columns (Cytiva). Samples were separated on a 0.6% alkaline agarose gel run at 20V for 16 hours in Alkaline Running Buffer (30 mM sodium hydroxide, 2 mM EDTA). Gels were dried onto Amersham Hybond-XL (GE Healthcare) and imaged using a phosphor screen, and scanned using an Amersham Typhoon phosphorimager (Cytiva).

## Supplementary Information

**Supplemental Figure 1.**
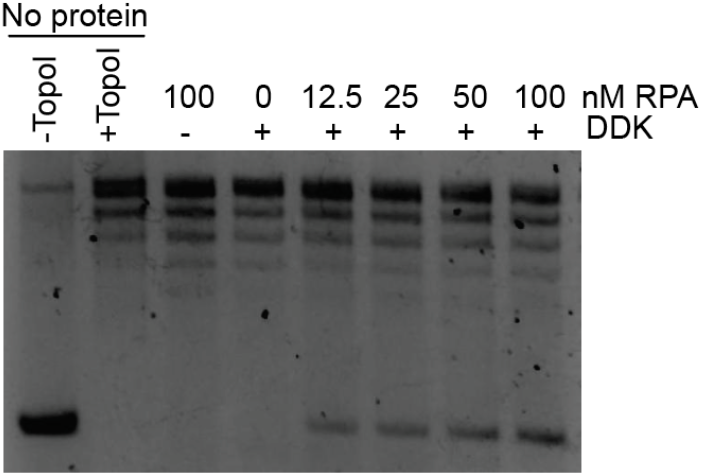
RPA titration in the plasmid unwinding assay shows that unwinding is both DDK-specific unwinding that requires sufficient concentrations of RPA.

**Supplemental Figure 2.**
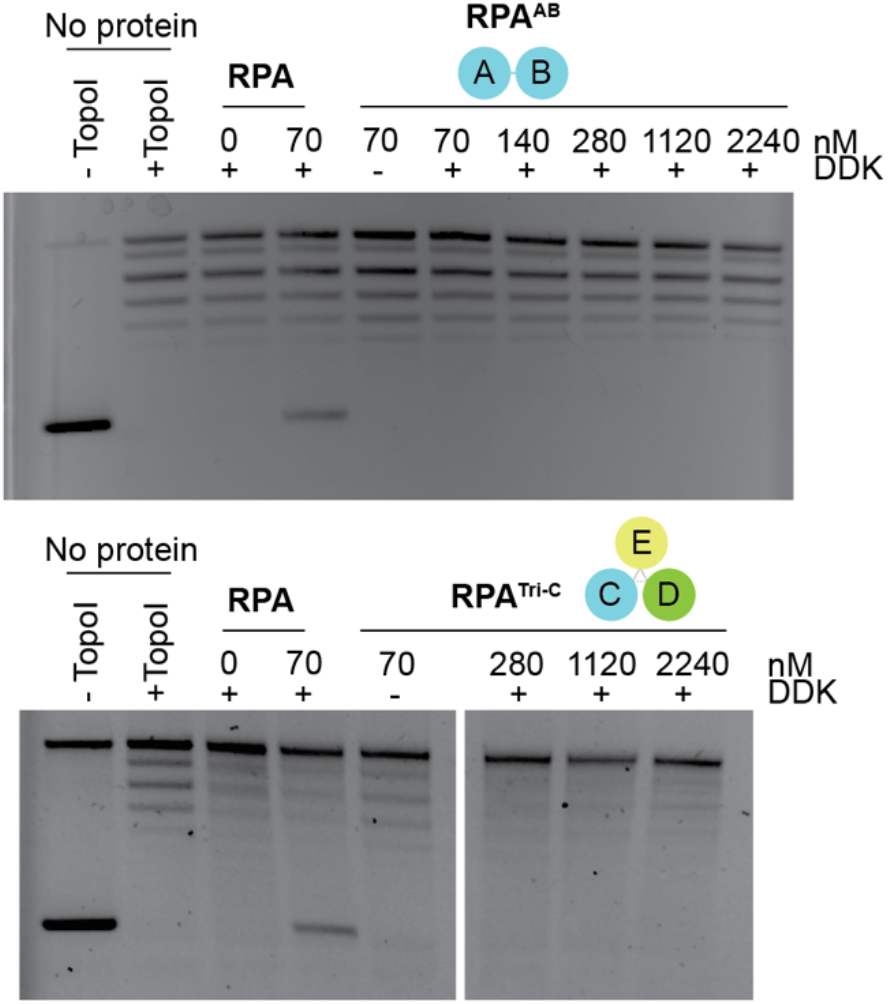
Extended titration series of RPA AB (top) and Tri-C (bottom) mutants going up to 2.24μM showed no observable DNA unwinding.

**Supplemental Figure 3.**
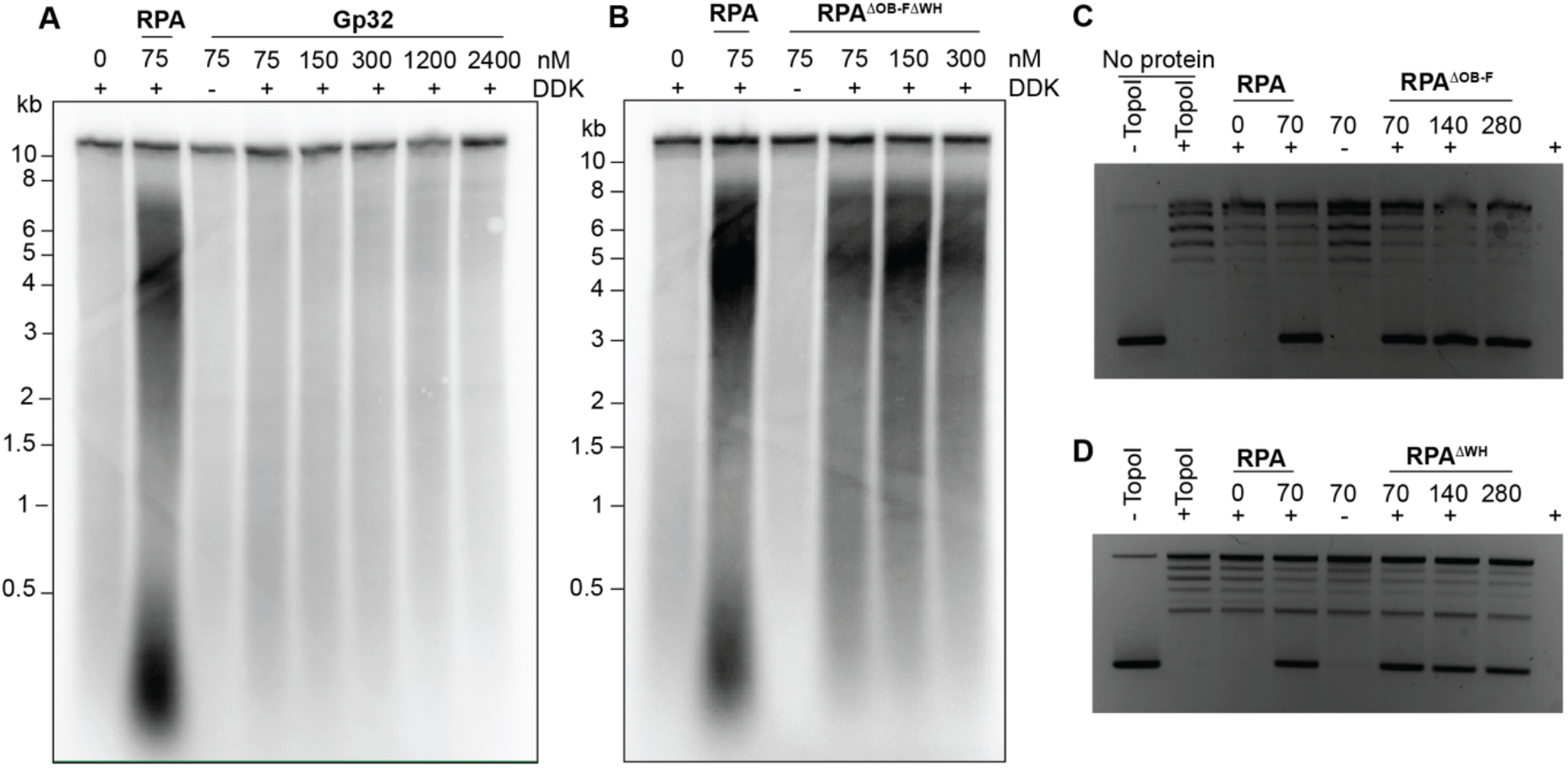
A) Gp32 does not support DNA replication in eukaryotic replication assays. B) Replication assay with RPA ΔOB-FΔWH. CD) Unwinding assays with RPA single domain deletions show that, like the double mutant, the single domain deletions support DNA unwinding.

**Supplemental Table 1.** Apparent Kd values and 95% confidence intervals from anisotropy data fit to the Hill Equation.

| Protein | K<sub>d,app</sub> | 95% CI | Figure |
| --- | --- | --- | --- |
| RPA | 1.729 | 1.480 to 2.015 | 1,3,4 |
| EcSSB | 0.114 | 0.08112 to 0.1484 | 1 |
| Gp32 | 18.670 | 15.25 to 23.03 | 1 |
| RPA-AB | 18.350 | 14.00 to 24.69 | 3 |
| RPA-TriC | 2.971 | 2.133 to 4.648 | 3 |
| RPA-ABAB | 1.450 | 1.182 to 1.794 | 4 |
| RPA-ΔOB-F,ΔWH | 2.047 | 1.402 to 3.496 | 4 |

**Supplemental Table 2.**
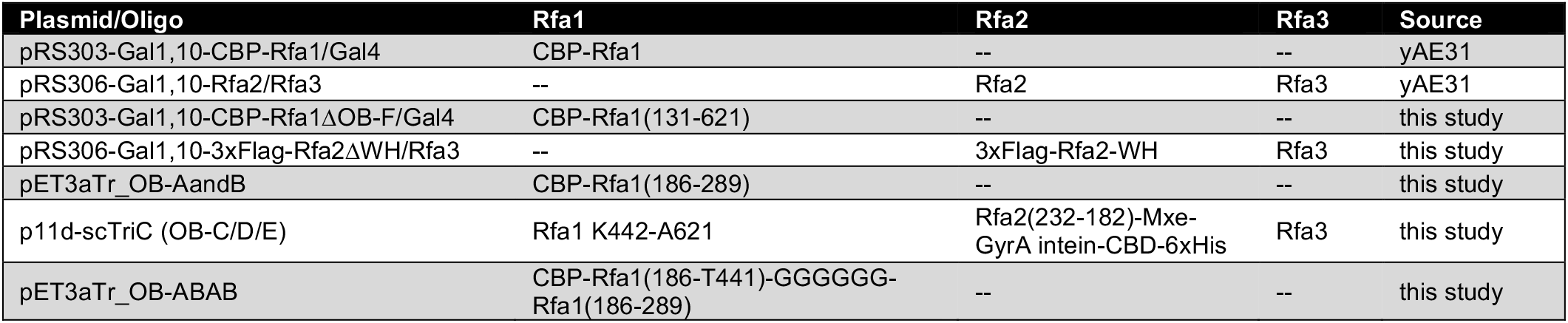
RPA Expression Plasmids.

**Supplemental Table 3.**
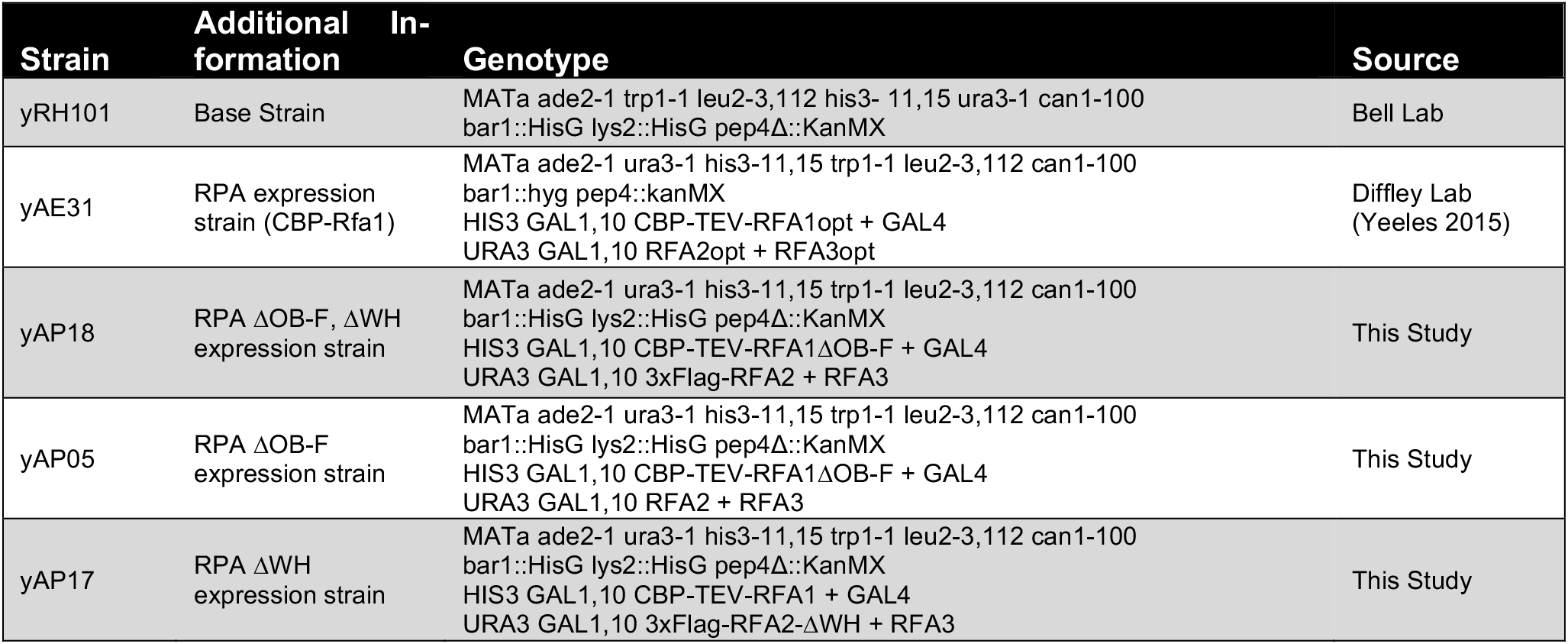
Yeast Strains.

## References

Acharya, A., Kasaciunaite, K., Göse, M., Kissling, V., Guérois, R., Seidel, R., and Cejka, P. (2021). Distinct RPA domains promote recruitment and the helicase-nuclease activities of Dna2. Nat. Commun. 12, 6521.

Ahmad, F., Patterson, A., Deveryshetty, J., Mattice, J.R., Pokhrel, N., Bothner, B., and Antony, E. (2021). Hydrogen-deuterium exchange reveals a dynamic DNA-binding map of replication protein A. Nucleic Acids Res. 49, 1455–1469.

Alani, E., Thresher, R., Griffith, J.D., and Kolodner, R.D. (1992). Characterization of DNA-binding and strand-exchange stimulation properties of y-RPA, a yeast single-strand-DNA-binding protein. J. Mol. Biol. 227, 54–71.

Bastin-Shanower, S.A., and Brill, S.J. (2001). Functional analysis of the four DNA binding domains of Replication Protein A: the role of RPA2 in ssDNA binding. J. Biol. Chem. 276, 36446–36453.

Bell, S.P., and Labib, K. (2016). Chromosome duplication in Saccharomyces cerevisiae. Genetics 203, 1027–1067.

Binz, S.K., Dickson, A.M., Haring, S.J., and Wold, M.S. (2006). Functional Assays for Replication Protein A (RPA). In Methods in Enzymology, pp. 11–38.

Bochkarev, A., Pfuetzner, R.A., Edwards, A.M., and Frappier, L. (1997). Structure of the single-stranded-DNA-binding domain of replication protein A bound to DNA. Nature 385, 176–181.

Bochkareva, E., Korolev, S., Lees-Miller, S.P., and Bochkarev, A. (2002). Structure of the RPA trimerization core and its role in the multistep DNA-binding mechanism of RPA. EMBO J. 21, 1855–1863.

Braun, K.A., Lao, Y., He, Z., Ingles, C.J., and Wold, M.S. (1997). Role of protein-protein interactions in the function of replication protein A (RPA): RPA modulates the activity of DNA polymerase alpha by multiple mechanisms. Biochemistry 36, 8443–8454.

Brill, S.J., and Stillman, B. (1989). Yeast replication factor-A functions in the unwinding of the SV40 origin of DNA replication. Nature 342, 92–95.

Brill, S.J., and Stillman, B. (1991). Replication factor-A from Saccharomyces cerevisiae is encoded by three essential genes coordinately expressed at S phase. Genes Dev. 5, 1589–1600.

Caldwell, C.C., and Spies, M. (2020). Dynamic elements of replication protein A at the crossroads of DNA replication, recombination, and repair. Crit. Rev. Biochem. Mol. Biol. 55, 482–507.

Collins, K.L., and Kelly, T.J. (1991). Effects of T antigen and replication protein A on the initiation of DNA synthesis by DNA polymerase alpha-primase. Mol. Cell. Biol. 11, 2108–2115.

van Deursen, F., Sengupta, S., De Piccoli, G., Sanchez-Diaz, A., and Labib, K. (2012). Mcm10 associates with the loaded DNA helicase at replication origins and defines a novel step in its activation. EMBO J. 31, 2195–2206.

Devbhandari, S., Jiang, J., Kumar, C., Whitehouse, I., and Remus, D. (2017). Chromatin Constrains the Initiation and Elongation of DNA Replication. Mol. Cell 65, 131–141.

Dornreiter, I., Erdile, L.F., Gilbert, I.U., von Winkler, D., Kelly, T.J., and Fanning, E. (1992). Interaction of DNA polymerase alpha-primase with cellular replication protein A and SV40 T antigen. EMBO J. 11, 769–776.

Douglas, M.E., Ali, F.A., Costa, A., and Diffley, J.F.X. (2018). The mechanism of eukaryotic CMG helicase activation. Nature 555, 265–268.

Eickhoff, P., Kose, H.B., Martino, F., Petojevic, T., Abid Ali, F., Locke, J., Tamberg, N., Nans, A., Berger, J.M., Botchan, M.R., et al. (2019). Molecular Basis for ATP-Hydrolysis-Driven DNA Translocation by the CMG Helicase of the Eukaryotic Replisome. Cell Rep. 28, 2673–2688.e8.

Evrin, C., Clarke, P., Zech, J., Lurz, R., Sun, J., Uhle, S., Li, H., Stillman, B., and Speck, C. (2009). A double-hexameric MCM2-7 complex is loaded onto origin DNA during licensing of eukaryotic DNA replication. Proc. Natl. Acad. Sci. 106, 20240–20245.

Fan, J., and Pavletich, N.P. (2012). Structure and conformational change of a replication protein A heterotrimer bound to ssDNA. Genes Dev. 26, 2337–2347.

Fanning, E., Klimovich, V., and Nager, A.R. (2006). A dynamic model for replication protein A (RPA) function in DNA processing pathways. Nucleic Acids Res. 34, 4126–4137.

Frigola, J., Remus, D., Mehanna, A., and Diffley, J.F.X. (2013). ATPase-dependent quality control of DNA replication origin licensing. Nature 495, 339–343.

Gambus, A., Khoudoli, G.A., Jones, R.C., and Blow, J.J. (2011). MCM2-7 form double hexamers at licensed origins in Xenopus egg extract. J. Biol. Chem. 286, 11855–11864.

Georgescu, R., Yuan, Z., Bai, L., De Luna Almeida Santos, R., Sun, J., Zhang, D., Yurieva, O., Li, H., and O’Donnell, M.E. (2017). Structure of eukaryotic CMG helicase at a replication fork and implications to replisome architecture and origin initiation. Proc. Natl. Acad. Sci. 114, E697–E706.

Gibb, B., Ye, L.F., Gergoudis, S.C., Kwon, Y.H., Niu, H., Sung, P., and Greene, E.C. (2014). Concentration-dependent exchange of replication protein A on single-stranded DNA revealed by single-molecule imaging. PLoS One 9.

Goswami, P., Abid Ali, F., Douglas, M.E., Locke, J., Purkiss, A., Janska, A., Eickhoff, P., Early, A., Nans, A., Cheung, A.M.C., et al. (2018). Structure of DNA-CMG-Pol epsilon elucidates the roles of the non-catalytic polymerase modules in the eukaryotic replisome. Nat. Commun. 9, 5061.

Han, Y., Loo, Y., Militello, K.T., and Melendy, T. (1999). Interactions of the Papovavirus DNA Replication Initiator Proteins, Bovine Papillomavirus Type 1 E1 and Simian Virus 40 Large T Antigen, with Human Replication Protein A. J. Virol. 73, 4899–4907.

Ilves, I., Petojevic, T., Pesavento, J.J., and Botchan, M.R. (2010). Activation of the MCM2-7 helicase by association with Cdc45 and GINS proteins. Mol. Cell 37, 247–258.

Ishimi, Y., Matsumoto, K., and Ohba, R. (1994). DNA replication from initiation zones of mammalian cells in a model system. Mol. Cell. Biol. 14, 6489–6496.

Jarmoskaite, I., AlSadhan, I., Vaidyanathan, P.P., and Herschlag, D. (2020). How to measure and evaluate binding affinities. Elife 9, 1–34.

Kang, S., Warner, M.D., and Bell, S.P. (2014). Multiple Functions for Mcm2-7 ATPase Motifs during Replication Initiation. Mol. Cell 55, 655–665.

Kenny, M.K., Lee, S.H., and Hurwitz, J. (1989). Multiple functions of human single-stranded-DNA binding protein in simian virus 40 DNA replication: single-strand stabilization and stimulation of DNA polymerases alpha and delta. Proc. Natl. Acad. Sci. 86, 9757–9761.

Kose, H.B., Xie, S., Cameron, G., Strycharska, M.S., and Yardimci, H. (2020). Duplex DNA engagement and RPA oppositely regulate the DNA-unwinding rate of CMG helicase. Nat. Commun. 11, 1–15.

Kowalczykowski, S.C., Lonberg, N., Newport, J.W., and von Hippel, P.H. (1981). Interactions of bacteriophage T4-coded gene 32 protein with nucleic acids. J. Mol. Biol. 145, 75–104.

Langston, L.D., and O’Donnell, M.E. (2019). An explanation for origin unwinding in eukaryotes. Elife 8, 1–17.

Lewis, J.S., Gross, M.H., Sousa, J., Henrikus, S.S., Greiwe, J.F., Nans, A., Diffley, J.F.X., and Costa, A. (2022). Mechanism of replication origin melting nucleated by CMG helicase assembly. Nature 606, 1007.

Li, H., and O’Donnell, M.E. (2018). The Eukaryotic CMG Helicase at the Replication Fork: Emerging Architecture Reveals an Unexpected Mechanism. BioEssays News Rev. Mol. Cell. Dev. Biol. 40.

Li, N., Zhai, Y., Zhang, Y., Li, W., Yang, M., Lei, J., Tye, B.K., and Gao, N. (2015). Structure of the eukaryotic MCM complex at 3.8 Å. Nat. 2015 5247564 524, 186–191.

Lohman, T. (1994). Escherichia coli Single-Stranded DNA-Binding Proteins: Multiple DNA-Binding Modes and Cooperativities. Annu. Rev. Biochem. 63, 527–570.

Lõoke, M., Maloney, M.F., and Bell, S.P. (2017). Mcm10 regulates DNA replication elongation by stimulating the CMG replicative helicase. Genes Dev. 31, 291–305.

Marceau, A.H. (2012). Functions of Single-Strand DNA-Binding Proteins in DNA Replication, Recombination, and Repair. In Single-Stranded DNA Binding Proteins, (Totowa, NJ: Humana Press), pp. 1–21.

Melendy, T., and Stillman, B. (1993). An interaction between replication protein A and SV40 T antigen appears essential for primosome assembly during SV40 DNA replication. J. Biol. Chem. 268, 3389–3395.

Mer, G., Bochkarev, A., Gupta, R., Bochkareva, E., Frappier, L., Ingles, C.J., Edwards, A.M., and Chazin, W.J. (2000). Structural Basis for the Recognition of DNA Repair Proteins UNG2, XPA, and RAD52 by Replication Factor RPA. Cell 103, 449–456.

Naufer, M.N., Morse, M., Möller, G.B., McIsaac, J., Rouzina, I., Beuning, P.J., and Williams, M.C. (2019). Self-regulation of single-stranded DNA wrapping dynamics by E. coli SSB promotes both stable binding and rapid dissociation. BioRxiv 2019.12.20.885368.

Naufer, M.N., Morse, M., Mcisaac, J., Rouzina, I., Beuning, P.J., and Williams, M.C. (2021). Multiprotein E. coli SSB - ssDNA complex shows both stable binding and rapid dissociation due to interprotein interactions. 49, 1532–1549.

Parker, M.W., Botchan, M.R., and Berger, J.M. (2017). Mechanisms and regulation of DNA replication initiation in eukaryotes. Crit. Rev. Biochem. Mol. Biol. 52, 107–144.

Pokhrel, N., Caldwell, C.C., Corless, E.I., Tillison, E.A., Tibbs, J., Jocic, N., Tabei, S.M.A., Wold, M.S., Spies, M., and Antony, E. (2019). Dynamics and selective remodeling of the DNA-binding domains of RPA. Nat. Struct. Mol. Biol. 26, 129–136.

Posse, V., Johansson, E., and Diffley, J.F.X. (2021). Eukaryotic DNA replication with purified budding yeast proteins (Elsevier Inc.).

Prakash, A., and Borgstahl, G.E.O. (2012). The Eukaryotic Replisome: a Guide to Protein Structure and Function.

Raghunathan, S., Kozlov, A.G., Lohman, T.M., and Waksman, G. (2000). Structure of the DNA binding domain of E. coli SSB bound to ssDNA. Nat. Struct. Biol. 7, 648–652.

Remus, D., Beuron, F., Tolun, G., Griffith, J.D., Morris, E.P., and Diffley, J.F.X. (2009). Concerted loading of Mcm2-7 double hexamers around DNA during DNA replication origin licensing. Cell 139, 719–730.

Rouzina, I., Pant, K., Karpel, R.L., and Williams, M.C. (2005). Theory of Electrostatically Regulated Binding of T4 Gene 32 Protein to Single- and Double-Stranded DNA. Biophys. J. 89, 1941–1956.

Shen, J., Zhao, Y., Pham, N.T., Li, Y., Zhang, Y., Trinidad, J., Ira, G., Qi, Z., and Niu, H. (2022). Deciphering the mechanism of processive ssDNA digestion by the Dna2-RPA ensemble. Nat. Commun. 13, 359.

Shorrocks, A.M.K., Jones, S.E., Tsukada, K., Morrow, C.A., Belblidia, Z., Shen, J., Vendrell, I., Fischer, R., Kessler, B.M., and Blackford, A.N. (2021). The Bloom syndrome complex senses RPA-coated single-stranded DNA to restart stalled replication forks. Nat. Commun. 12, 1–15.

Tsurimoto, T., and Stillman, B. (1991). Replication factors required for SV40 DNA replication in vitro. II. Switching of DNA polymerase alpha and delta during initiation of leading and lagging strand synthesis. J. Biol. Chem. 266, 1961–1968.

Waga, S., and Stillman, B. (1998). The DNA replication fork in eukaryotic cells. Annu. Rev. Biochem. 67, 721–751.

Weisshart, K., Taneja, P., and Fanning, E. (1998). The Replication Protein A Binding Site in Simian Virus 40 (SV40) T Antigen and Its Role in the Initial Steps of SV40 DNA Replication. J. Virol. 72, 9771–9781.

Wobbe, C.R., Weissbach, L., Borowiec, J.A., Dean, F.B., Murakami, Y., Bullock, P., and Hurwitz, J. (1987). Replication of simian virus 40 origin-containing DNA in vitro with purified proteins. Proc. Natl. Acad. Sci. 84,1834-1838.

Wold, M.S. (1997). Replication protein A: a heterotrimeric, single-stranded DNA-binding protein required for eukaryotic DNA metabolism. Annu. Rev. Biochem. 66, 61–92.

Wold, M.S., and Kelly, T. (1988). Purification and characterization of replication protein A, a cellular protein required for in vitro replication of simian virus 40 DNA. Proc. Natl. Acad. Sci. U. S. A. 85, 2523–2527.

Wold, M.S., Weinberg, D.H., Virshup, D.M., Li, J.J., and Kelly, T.J. (1989). Identification of cellular proteins required for simian virus 40 DNA replication. J. Biol. Chem. 264, 2801–2809.

Yao, N.Y., and O’Donnell, M.E. (2021). The DNA Replication Machine: Structure and Dynamic Function. Subcell. Biochem. 96, 233–258.

Yates, L.A., Aramayo, R.J., Pokhrel, N., Caldwell, C.C., Kaplan, J.A., Perera, R.L., Spies, M., Antony, E., and Zhang, X. (2018). A structural and dynamic model for the assembly of Replication Protein A on single-stranded DNA. Nat. Commun. 9, 5447.

Yeeles, J.T.P., Deegan, T.D., Janska, A., Early, A., and Diffley, J.F.X. (2015). Regulated eukaryotic DNA replication origin firing with purified proteins. Nature 519, 431–435.

Yuzhakov, A., Kelman, Z., Hurwitz, J., and O’Donnell, M. (1999). Multiple competition reactions for RPA order the assembly of the DNA polymerase delta holoenzyme. EMBO J. 18, 6189–6199.

Zou, L., and Elledge, S.J. (2003). Sensing DNA Damage Through ATRIP Recognition of RPA-ssDNA Complexes. Science (80-.). 300, 1542–1548.

